# Distinct structure-function relationships across cortical regions and connectivity scales in the rat brain

**DOI:** 10.1101/742833

**Authors:** Milou Straathof, Michel R.T. Sinke, Theresia J.M. Roelofs, Erwin L.A. Blezer, R. Angela Sarabdjitsingh, Annette van der Toorn, Oliver Schmitt, Willem M. Otte, Rick M. Dijkhuizen

## Abstract

An improved understanding of the structure-function relationship in the brain is necessary to know to what degree structural connectivity underpins abnormal functional connectivity seen in many disorders. We integrated high-field resting-state fMRI-based functional connectivity with high-resolution macro-scale diffusion-based and meso-scale neuronal tracer-based structural connectivity, to obtain an accurate depiction of the structure-function relationship in the rat brain. Our main goal was to identify to what extent structural and functional connectivity strengths are correlated, macro- and meso-scopically, across the cortex. Correlation analyses revealed a positive correspondence between functional connectivity and macro-scale diffusion-based structural connectivity, but no correspondence between functional connectivity and meso-scale neuronal tracer-based structural connectivity. Locally, strong functional connectivity was found in two well-known resting-state networks: the sensorimotor and default mode network. Strong functional connectivity within these networks coincided with strong short-range intrahemispheric structural connectivity, but with weak heterotopic interhemispheric and long-range intrahemispheric structural connectivity. Our study indicates the importance of combining measures of connectivity at distinct hierarchical levels to accurately determine connectivity across networks in the healthy and diseased brain. Distinct structure-function relationships across the brain can explain the organization of networks and may underlie variations in the impact of structural damage on functional networks and behavior.

---

The brain is a complex organ that can be regarded as a structural and functional network of interacting regions at the micro-, meso- and macroscopic level. At the macro-scale, whole-brain functional networks can be non-invasively mapped with resting-state functional MRI (resting-state fMRI). In resting-state fMRI data the inter-regional temporal correlations of spontaneous low-frequency blood oxygenation level-dependent (BOLD) fluctuations reflect functional connectivity (Biswal et al. 1995; Fox and Raichle 2007). Based on clusters of functionally connected regions, various resting-state networks have been identified, such as the default mode network (Greicius et al. 2003). These networks have been related to behavioral functioning in health and disease, and abnormalities partially explain pathophysiological processes and disease progression (van den Heuvel and Hulshoff Pol 2010; Zhang and Raichle 2010; Raichle 2015).

The exact nature of functional connectivity is nonetheless not yet fully established. Since functional connectivity measured with resting-state fMRI relies on synchronous BOLD signals, understanding functional connectivity starts with understanding the origin of BOLD signals. The BOLD signal captures hemodynamic changes, such as blood flow, in response to neural activity. Although it is clear that BOLD signals reflect aspects of neural responses, it is still unclear which processes are the main contributors, i.e. excitation or inhibition, local field potentials, action potentials or multi-unit activity (Logothetis et al. 2001; Logothetis and Wandell 2004; Logothetis 2008). We know from primate and rodent studies that spontaneous BOLD fluctuations match with slow fluctuations in neuronal activity (Shmuel and Leopold 2008; Pan et al. 2011; Magri et al. 2012). Moreover, in humans, BOLD signal fluctuations are related to slow cortical potentials and gamma band-limited power (He et al. 2008). Still, the underlying structure of functional connectivity remains largely unknown. Since functional connectivity is found between adjacent and remote brain areas, short- and long-distance structural connections seem essential. Structural connectivity can be measured non-invasively with diffusion MRI and invasively with neuronal tracers. Diffusion-based tractography enables reconstruction of whole-brain macro-scale structural networks, by indirectly inferring the direction and strength of large white matter tracts from the diffusion of water (Turner et al. 1990; Basser et al. 1994). In contrast, neuronal tracers use the transport mechanisms of cells to label existing mono- or polysynaptic connections. Tracers thus provide a direct and accurate measure of the directionality and strength at the meso-scale of individual axonal projections (Heimer and Robards 1981).

Functional connectivity strength correlates with both diffusion- and neuronal tracer-based structural connectivity strength at the whole-brain level (Honey et al. 2009; Miranda-Dominguez et al. 2014); for an overview see (Straathof et al. 2019). However, different regions and connections display different structure-function relationships (Wang et al. 2012; Zimmermann et al. 2016; Grandjean et al. 2017). Identifying where and to what extent structural and functional connectivity strengths correlate will help to understand how brain networks are organized, and why functional abnormalities in brain disorders are related to characteristic patterns of disconnection or reorganization. So far, most studies have compared functional connectivity with structural connectivity measured at either the macro-scale or meso-scale, and thereby did not capture all aspects of structural connectivity. In addition, studies that applied diffusion MRI are hampered by the fact that a diffusion-based structural network is a suboptimal reconstruction of macro-scale axonal projections (Thomas et al. 2014; Maier-Hein et al. 2017; Schilling et al. 2018; Sinke et al. 2018). More accurate assessment of the structure-function relationships requires integration of functional connectivity with both macro-scale diffusion- and meso-scale neuronal tracer-based structural measures. Distinct structure-function relationships may be present at these different hierarchical levels (Reid et al. 2016). Rodents are excellent species to study these relationships as resting-state fMRI and diffusion MRI-based tractography are feasible in rodents (Dijkhuizen and Nicolay 2003) and comprehensive rodent databases of neuronal tracer-based structural connectivity are available as well (Schmitt and Eipert 2012; Noori et al. 2017).

In this study we combined high-field resting-state fMRI-based functional connectivity measurements and diffusion-as well as neuronal tracer-based structural connectivity measurements from the rat brain to spatially map the structure-function relationship at the macro- and meso-scale. Our main goal was to identify to what extent structural and functional connectivity strength are correlated, macro- and meso-scopically, across the rat brain, which could explain differences in the functional significance of connections and their contribution to network dysfunction in brain disorders. We distinguished interhemispheric and intrahemispheric connections, as well as specific functional networks (sensorimotor or default mode network).

## Materials and Methods

### Ethics statement

All experiments were approved by the Committee for Animal Experiments of the University Medical Center Utrecht, The Netherlands, and were conducted in agreement with European regulations (Guideline 86/609/EEC) and Dutch laws (‘Wet op de Dierproeven’, 1996).

### Animals

#### In vivo resting-state functional connectivity

Resting-state functional connectivity was measured in twelve healthy adult male Wistar rats with a weight of 479 ± 44 g (mean ± standard deviation (SD)), which were group-housed and used for an earlier described study (Roelofs et al. 2017). All animals had *ad libitum* access to food and water and were housed under the same environmental conditions (temperature 22-24° and 12 h light/dark cycle with lights on at 7:00 AM).

#### Post-mortem diffusion-based structural connectivity

Diffusion-based structural connectivity was measured in ten healthy adult male Wistar rats with an age of around twelve weeks. These animals were previously used in another study (Sarabdjitsingh et al. 2017) and group-housed under standard environmental conditions (12h light/dark cycle with lights on at 7:00 AM). Animals were sacrificed by an intraperitoneal pentobarbital injection followed by transcardial perfusion-fixation with 4% paraformaldehyde in phosphate-buffered saline, as previously described (Sarabdjitsingh et al. 2017). We extracted the brains, while leaving them inside the skull, and placed these in a proton-free oil (Fomblin®) prior to MR imaging to minimize susceptibility artefacts.

### MRI acquisition

All MRI experiments were conducted on a 9.4T horizontal bore Varian MR system (Palo Alto, CA, USA), equipped with a 400 mT/m gradient coil (Agilent).

#### In vivo resting-state functional connectivity

Before MRI, the animals were anesthetized (with 4% of isoflurane in air for induction). Endotracheal intubation was performed to mechanically ventilate the rats with 1.5% of isoflurane in a mixture of air and O2 (4:1). End-tidal CO2 was continuously monitored with a capnograph (Microcap, Oridion Medical 1987 Ltd., Jerusalem, Israel). The animals were placed in an animal cradle and immobilized in a specially designed stereotactic holder. During MRI, a feed-back controlled heating pad ensured that the body temperature of the rats was maintained at 37.0 ± 1.0 °C. Blood oxygen saturation and heart rate were monitored with a pulse-oximeter from signals recorded with an infrared sensor attached to the hind paw of the animal.

We used a home-built 90 mm diameter Helmholtz volume coil for radiofrequency transmission, and an inductively coupled 25 mm diameter surface coil for signal detection. Prior to resting-state fMRI acquisition we acquired an anatomical image for registration purposes using 3D balanced steady-state free precession (bSSFP) imaging with four phase-cycling angles (0°, 90°, 180°, 270°). The acquisition parameters were as follows: repetition time (TR) / echo time (TE) = 5/2.5 ms; flip angle = 20°; field-of-view (FOV) = 40×32×24 mm^3^; acquisition matrix = 160×128×96; image resolution = 250 μm isotropic. Total acquisition time = 12.5 min. Resting-state fMRI images were acquired with T_2_^*^-weighted blood oxygenation level-dependent (BOLD) single shot 3D gradient-echo Echo Planar Imaging (EPI). The acquisition parameters were as follows: TR/TE = 26.1/15 ms; flip angle = 13°; FOV = 32.4×32.4×16.8 mm^3^, Acquisition matrix = 54×54×28; Spatial Resolution = 600 μm isotropic. The acquisition time was 730.8 ms per volume, with a total of 800 volumes, resulting in a scan time of 9 minutes and 45 seconds.

#### Post-mortem diffusion-based structural connectivity

For diffusion MRI we used a custom-made solenoid coil with an internal diameter of 26 mm. High spatial and angular resolution diffusion imaging (HARDI) was performed with an 8-shot 3D EPI sequence. The acquisition parameters were as follows: TR/TE = 500/32.4 ms, Δ/δ = 15/4 ms; b-value = 3842 s/mm^2^; FOV = 19.2×16.2×33 mm^3^; Acquisition matrix = 128×108×220; Spatial resolution: 150×150×150 μm^3^. Diffusion-weighting was executed in 60 non-collinear directions on a half sphere and included five b_0_ non-diffusion-weighted images, with a total scan time of 8 hours.

### MRI processing

All MRI analyses were performed using FMRIB’s Software Library (FSL) v5.0, unless otherwise stated.

#### In vivo resting-state functional connectivity

The first twenty images of the resting-state fMRI scan were removed to ensure a steady state and the remaining images were motion-corrected to the mean volume with *MCFLIRT* (Jenkinson et al. 2002) and brain-extracted with *BET* (Smith 2002). The six motion correction parameters were used as regressors for the resting-state fMRI signal. No global signal regression was performed. Low-frequency BOLD fluctuations were obtained by band-pass filtering between 0.01 and 0.1 Hz in *AFNI* (Cox 1996). Fisher’s Z-transformed full correlation coefficients were calculated between the timeseries for all pairs of regions of interest (see below). These Fisher’s Z-transformed correlation coefficients were averaged over all rats within each dataset to obtain a group-level measurement of functional connectivity strength between our regions of interest.

#### Post-mortem diffusion-based structural connectivity

We used single shell constrained spherical deconvolution (CSD) to construct a fiber orientation distribution (FOD) map for every rat. Next, CSD-based tractography, using the iFOD2 algorithm, was performed in MRtrix3^®^ (http://www.mrtrix.org/) (Tournier et al. 2010, 2012). The iFOD2 algorithm uses 2^nd^ order integration over adjacent orientation distributions (Tournier et al. 2010). Whole brain tractography was done in individual subject space using dynamic seeding, thereby generating 2.5 million streamlines with a step size of 75 μm, an angle threshold of 40° and a FOD threshold of 0.2. The generated tractograms were filtered by *Spherical deconvolution Informed Filtering of Tracts* (SIFT) (Smith et al. 2013, 2015). Subsequently, the connectomes were constructed by matching the filtered tractograms with a custom-built 3D model of the 5^th^ edition of the Paxinos and Watson rat brain atlas (Paxinos and Watson 2005; Majka et al. 2012) in subject space. Regions of interest (see below) were structurally connected if one or multiple streamlines had their endpoints in both regions, where the filtered number of inter-regional streamlines was indicative of structural connectivity strength. Finally, we calculated an average weighted connectome, to obtain a group-level measurement of diffusion-based structural connectivity strength between our regions of interest.

#### Regions of interest

To enable the selection of regions of interest, the mean resting-state fMRI image of each dataset was first linearly registered (*FLIRT* (Jenkinson and Smith 2001; Jenkinson et al. 2002)) to the anatomical image of the same animal, followed by non-linear registration (*FNIRT* (Andersson et al. 2007)) to a custom-built 3D model of the Paxinos and Watson rat brain atlas (Paxinos and Watson 2005; Majka et al. 2012). For diffusion MRI, the average of the non-diffusion-weighted images of each individual rat was non-linearly registered to this rat brain atlas. These registrations were used to transform 106 cortical bilateral regions into individual diffusion MRI and resting-state fMRI spaces. We only included regions of interest with sufficient assurance of spatial alignment, i.e. regions consisting of at least 8 voxels in individual resting-state fMRI space. This resulted in 82 bilateral cortical regions (Table 1).

**Table 1:**
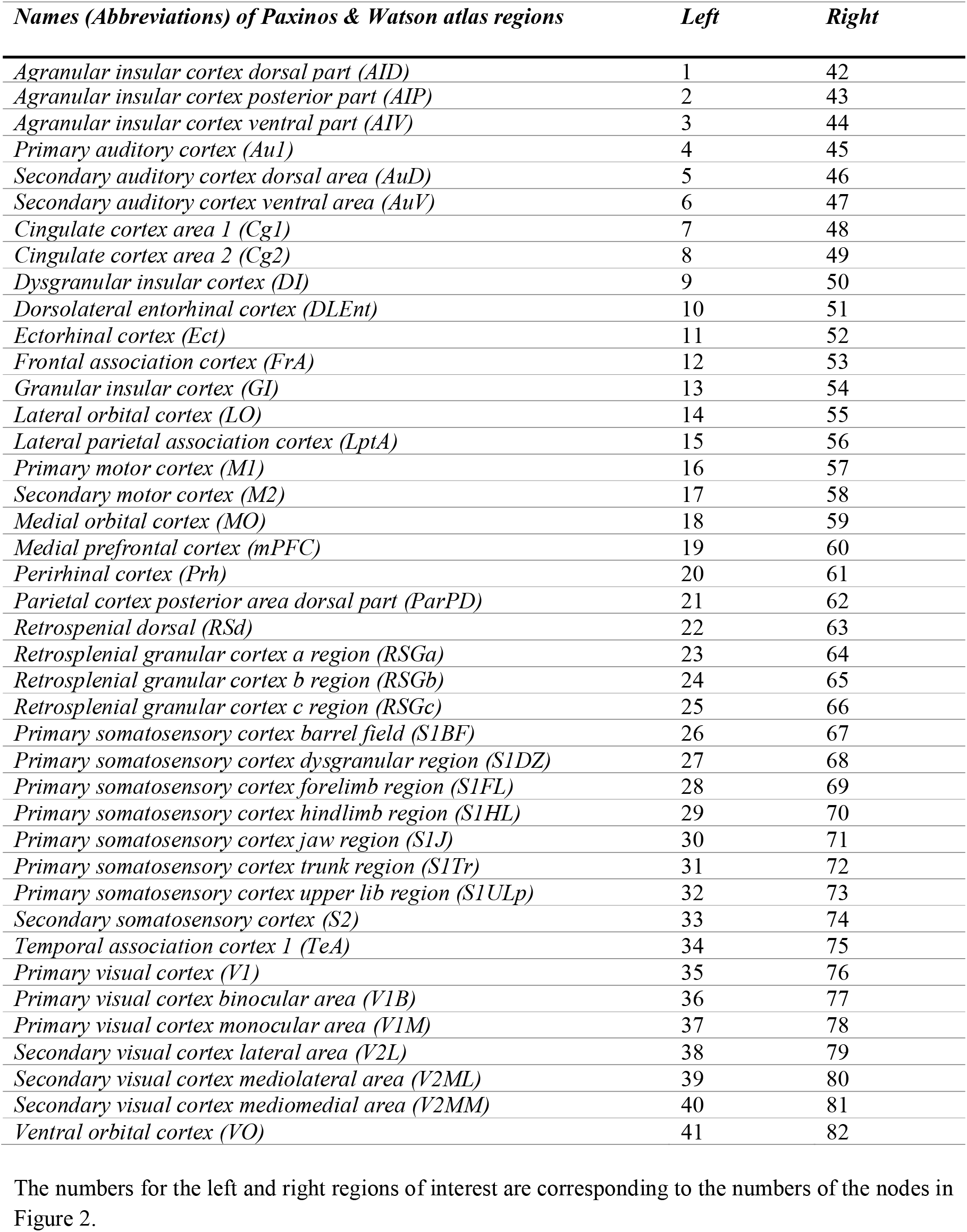
Included regions of interest for resting-state fMRI, diffusion MRI and neuronal tracer analyses.

### Neuronal tracer-based structural connectivity

Neuronal tracer-based structural connectivity data was extracted from the NeuroVIISAS database (Schmitt and Eipert 2012). This database contains rat nervous system data from over 7860 published tract-tracing studies, describing in total 591,435 ipsi- and contralateral connections. Many of these connections are described in multiple studies, affirming the robustness of the dataset. Studies with anterograde as well as retrograde monosynaptic tracers have been included, giving directionality information about the structural connections.

We used the same regions of interest as described for the functional connectivity and diffusion-based structural connectivity analyses to extract neuronal tracer-based structural connectivity for all pairs of regions (Supplementary table 1). The directional weight for each connection is assigned in the NeuroVIISAS database as follows: 0: no information available; 1: light/sparse connection; 2: moderate/dense connection; 3: strong connection and 4: very strong connection. These categorical descriptors were transformed to continuous data by using a logarithmic transformation. We averaged these continuous connection weights over all studies investigating the same connection, resulting in a continuous scale for neuronal tracer-based structural connectivity between 0 and 4.

### Experimental design and statistical analysis

The network of 82 regions consisted of 6,724 connections, of which we removed the self-connections, resulting in a total of 6,642 connections. Only connections that existed in both the macro- and mesoscale structural connectivity datasets, meaning that they had a structural connectivity strength higher than zero in both datasets, were included for the analysis, to exclude false-positives often present in diffusion-based tractography networks (Maier-Hein et al. 2017).

#### Relationship between structural and functional connectivity strength at whole-brain level

To map the structure-function relationship globally, we performed correlation analyses between functional connectivity strength and macro-scale diffusion-based or meso-scale neuronal tracer-based structural connectivity separately. We applied a log-transformation to both structural connectivity weights because they were skewed towards smaller connectivity weight values. We calculated a Spearman rank correlation coefficient (ρ) between functional connectivity strength and macro-scale diffusion-based or meso-scale neuronal tracer-based structural connectivity strength.

#### Relationship between structural and functional connectivity strength at connection level

To map the level of agreement between structural and functional connectivity at connection level, we selected the strongest and weakest structural connections at both the macro- and meso-scale. The strongest structural network was defined by connections that belonged to the 25% strongest diffusion-based and 25% strongest neuronal tracer-based structural connections. The weakest structural network was defined by connections that belonged to the 25% weakest diffusion-based and 25% weakest neuronal tracer-based structural connections. By combining macro- and meso-scale structural connectivity strengths, we selected the structural networks that were strong or weak at the level of individual axonal projections as well as at the level of large white matter bundles. This heightened the reliability of our assessment of the strength of structural connections and reduced the influence of methodological bias for specific connections.

For both the strongest and weakest structural network, we determined the 25% strongest and 25% weakest functional connections, resulting in four sub-groups of connections. Two of these subgroups represent connections where structural and functional connectivity strength agree: strong structural and functional connectivity or weak structural and functional connectivity. The other two subgroups are connections where structural and functional connectivity strength disagree: strong structural connectivity but weak functional connectivity or weak structural connectivity but strong functional connectivity.

To determine whether these subgroups of connections share common characteristics, we determined the Euclidian distance and type of connections and regions for all connections.

Between each pair of regions, we calculated the Euclidian distance, which is the shortest distance between two points in space (i.e. in a straight line). Therefore, we determined the x, y and z coordinate in mm of the center of gravity of each region in atlas space. Subsequently, we calculated the Euclidean distance, between each pair of regions i and j, with the following formula:

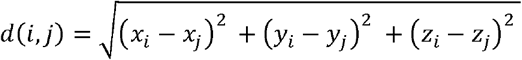

We divided the included connections and regions in sub-groups based on two different criteria. First, for each connection, we identified whether it was an intrahemispheric connection, which runs between two regions in the same hemisphere, or an interhemispheric connection, which runs between two regions in different hemispheres. In addition, we subdivided the interhemispheric connections into homotopic interhemispheric connections, which run between two homologous areas in different hemispheres and heterotopic interhemispheric connections, which run between two dissimilar areas in different hemispheres (Figure 1C).

**Figure 1:**
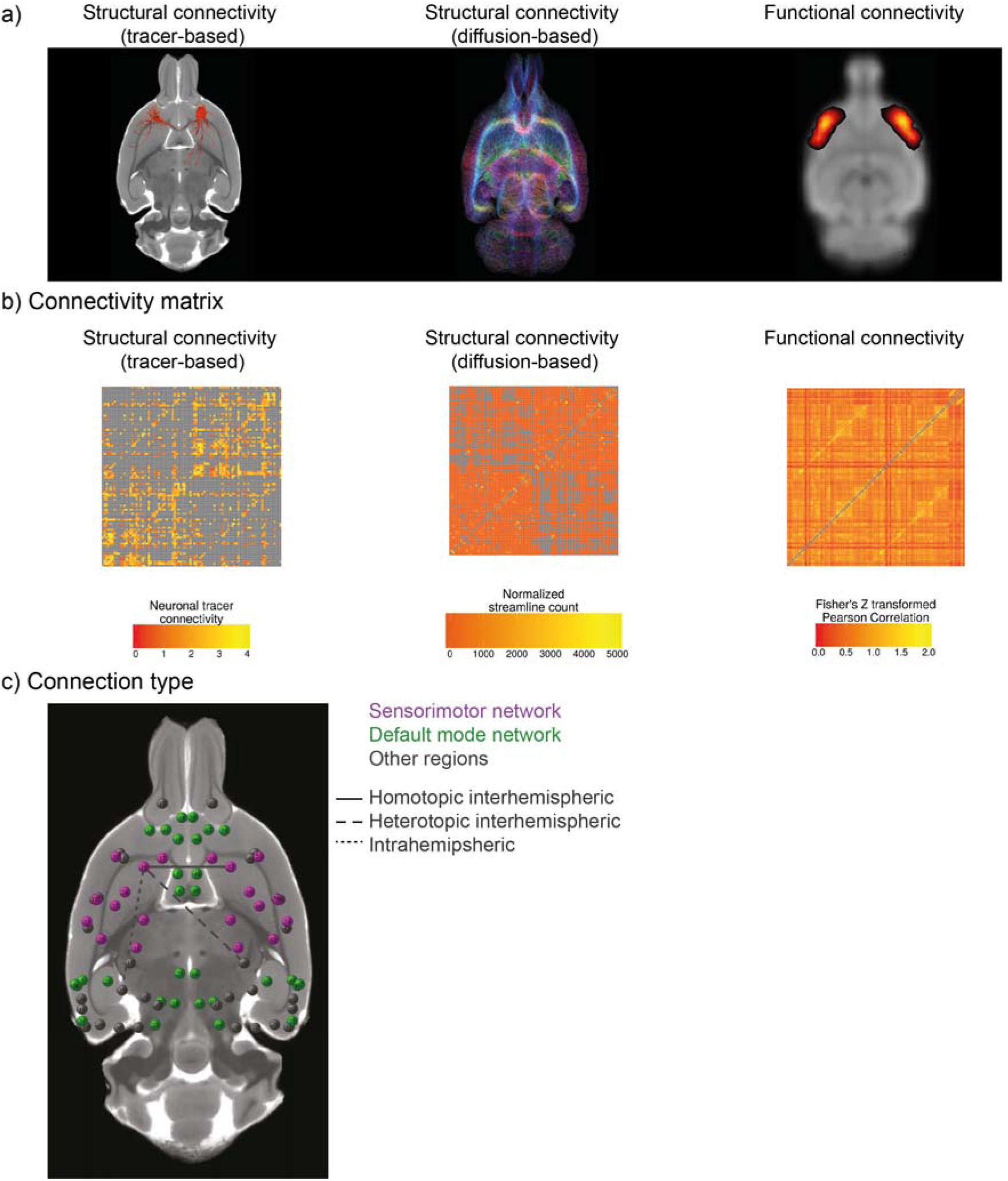
Overview of the analysis pipeline. Rat brain images are shown as axial views. Different measures of connectivity in the rat brain were assessed (a): meso-scale neuronal tracer-based structural connectivity (left), macro-scale diffusion-based structural connectivity (middle) and macroscale resting-state functional connectivity (right). For each measure, we determined the connectivity matrix between 82 cortical regions of interest, with exclusion of the self-connections (central diagonal line) (b). We combined all connectivity matrices to determine the structure-function connectome of the rat brain (c) (circles representing nodes). The connectomes are visualized in 3D. The colors in (c) represent two well-described functional resting-state networks in the rat brain: the sensorimotor network (purple) and the default mode network (green). Regions not belonging to these networks are shown in gray. The lines represent different regional types of connections: homotopic interhemispheric connection (solid line), heterotopic interhemispheric connection (dashed line) or intrahemispheric connection (dotted line).

Second, for each region of interest, we assessed whether it belonged to one of two well-described functional networks in the rat brain: the sensorimotor network or the default mode network (Figure 1C). The sensorimotor network was defined as consisting of the left and right primary and secondary motor cortex (M1 and M2), subdivisions of the primary somatosensory cortex (S1BF, S1DZ, S1FL, S1HL, S1J, S1Tr, S1ULp) and the secondary somatosensory cortex (S2) (Sierakowiak et al. 2015). The default mode network was defined as consisting of the left and right medial prefrontal cortex (mPFC), the cingulate cortex (Cg1 and Cg2), the orbital cortex (VO, MO and LO), the auditory/temporal association cortex (Au1, AuD, AuV and TeA), the posterior parietal cortex (ParPD) and the retrosplenial cortex (RSd, RSGb, RSGc) (Lu et al. 2012). For each connection, we determined whether the connection was within one of these functional networks, or whether it was connecting one of these functional networks with another functional network.

The analysis pipeline is illustrated in Figure 1. All statistical and descriptive analyses were performed in R (version 3.2.3) (R Core Team 2014).

## Results

Of all the possible 6,642 connections between the 82 selected regions of the cortical network, 1,175 connections (17.7% of total connections) displayed structural connectivity in both the diffusion MRI and the neuronal tracer dataset. The average Euclidean distance for all the included connections in this network was 6.08 ± 3.35 mm (mean ± standard deviation (SD)).

### Different pathways and brain circuits display distinct structure-function relationships

The strongest structural network at the macro- and meso-scale consisted of 107 cortical connections. These strongest structural connections were mainly intrahemispheric (93% of total connections; left: 47%, right: 46 %) with an average Euclidean distance of 2.54 ± 1.71 mm (see Figure 2). The weakest structural network at the macro- and meso-scale consisted of 93 connections. Of these weakest structural connections 31% was interhemispheric and 69% was intrahemispheric (left: 32%, right: 37%), with an average distance of 9.55 ± 2.42 mm (see Figure 2).

**Figure 2:**
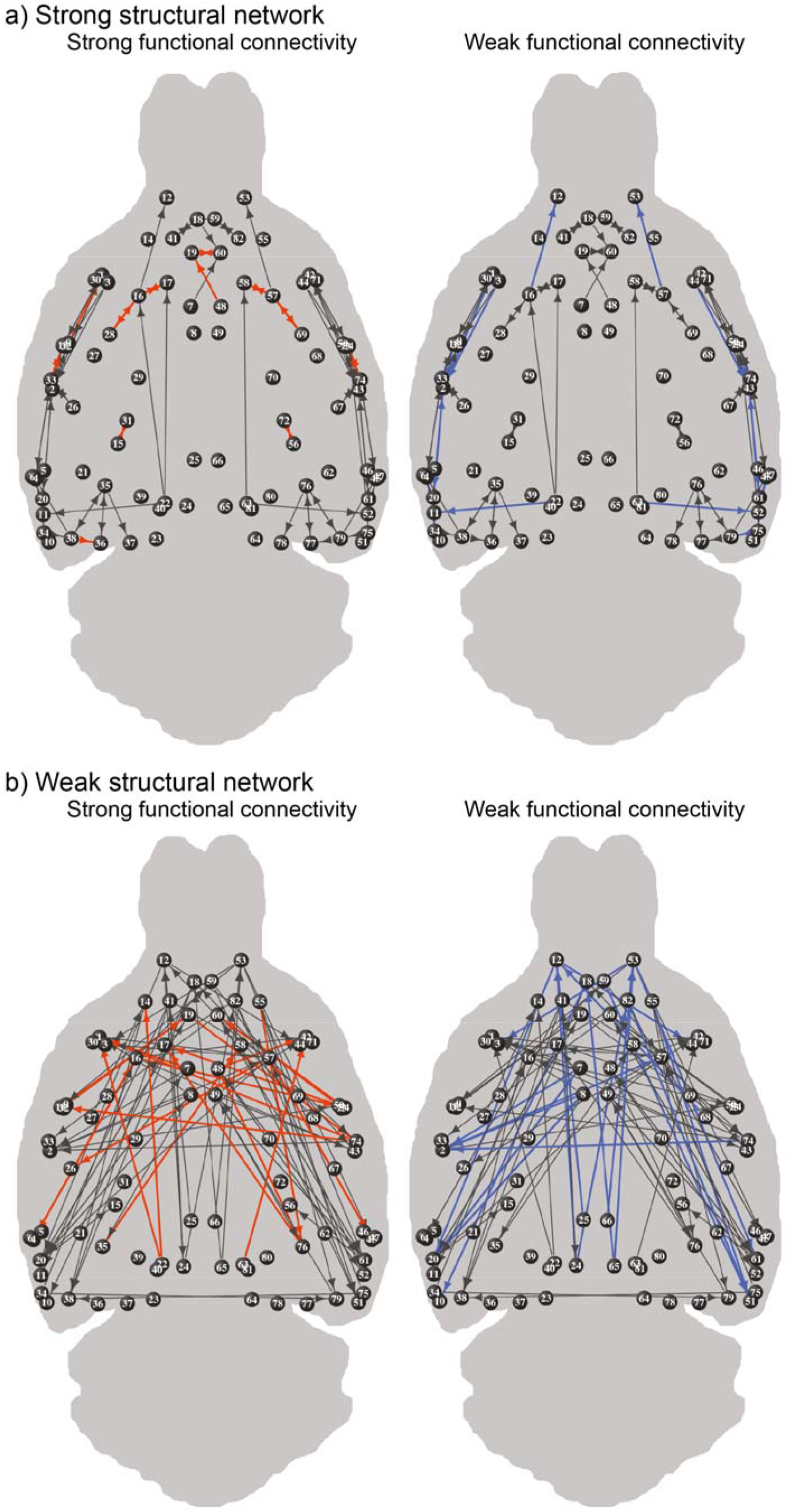
Strongest and weakest functional connections within the strongest and weakest structural networks. The strongest structural network consists of the 25% strongest structural connections at the macro-scale and meso-scale (a), and the weakest structural network consists of the 25% weakest (b). Functional connectivity is colored for the 25% strongest (red) and 25% weakest functional connections (blue). Circles represent the nodes (regions of interest), with numbers representing the regions listed in Table 1, and lines represent the edges (connections). The connectomes are visualized in 3D. The arrowheads reflect directionality information determined from the neuronal tracer-based structural connectivity dataset.

Within both the strongest and weakest cortical structural networks, we determined the 25% strongest and 25% weakest functional connections. These strongest and weakest functional connections are depicted in Figure 2. The characteristics of these subcategories of connections are summarized in Figure 3 and described below.

**Figure 3:**
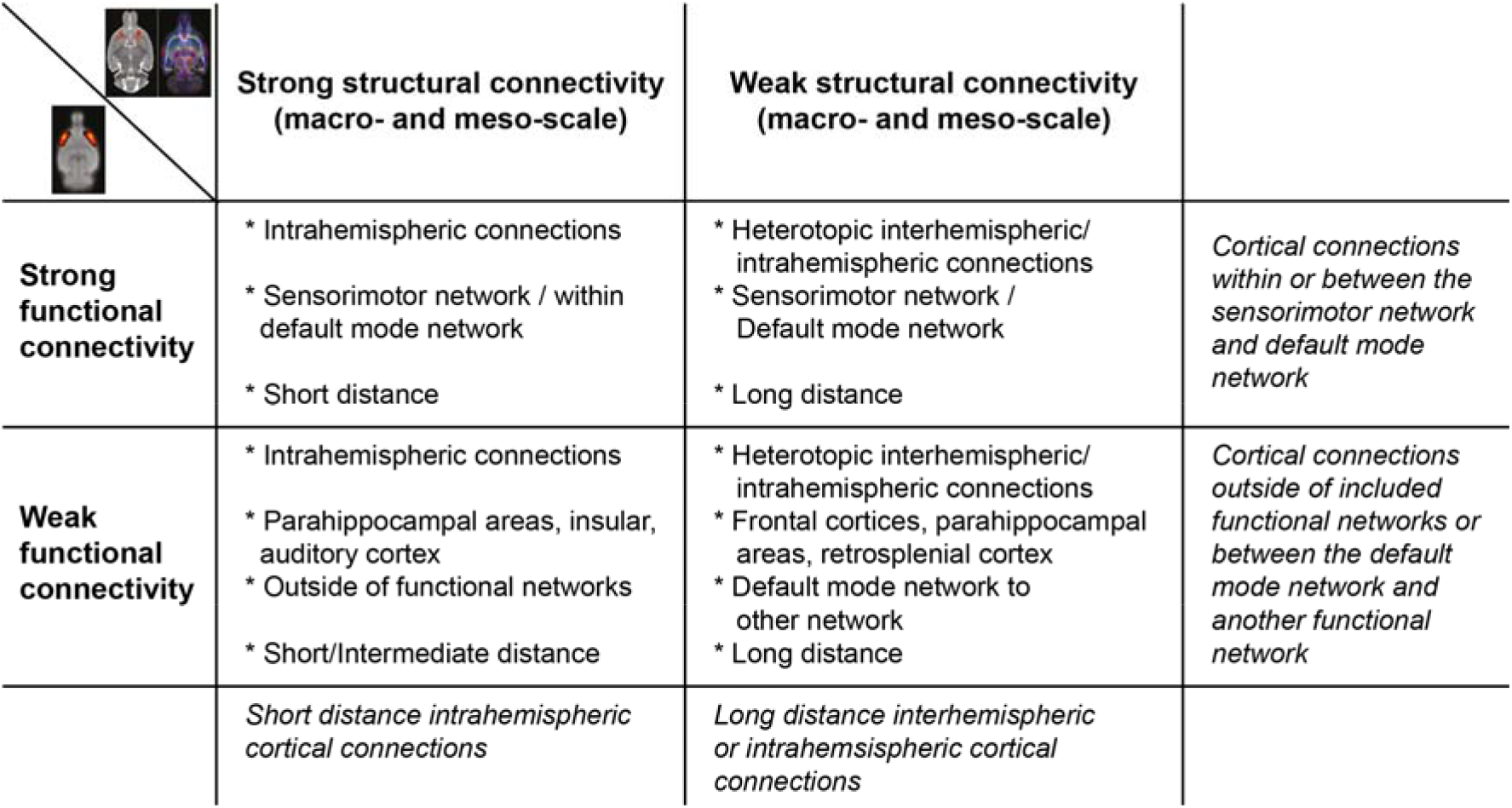
Characteristics of connections per subcategory of structural and functional connectivity. Structural connectivity is depicted in columns, whereas functional connectivity is depicted in rows. Strong connections belong to the 25% strongest connections; structurally based on diffusion MRI and neuronal tracing (i.e. at both the macro- and meso-scale) and functionally based on resting-state fMRI. Similarly, weak structural connections belong to the 25% weakest connections.

Connections with strong structural and functional connectivity are shown in Table 2. The average length of the connections was 1.34 ± 0.69 mm. Eighty-eight percent of these strongest connections was intrahemispheric (left: 50%; right: 38%). a Sixty-two percent of the connections were part of the sensorimotor network. The homotopic connection between the left and right medial prefrontal cortex, which is part of the default mode network, was also one of the identified strongest connections.

**Table 2:** Characteristics of cortical connections in the rat brain with strong meso- and macroscale structural connectivity and strong functional connectivity.

| Seed | Target | Neuronal<br>tracer-based<br>structural<br>connectivity<br>strength | Diffusion-based<br>structural<br>connectivity<br>strength | Functional<br>connectivity<br>strength (Z') | Euclidean<br>distance (mm) | Connection type<br>(network) | Connection type (regional) |
| --- | --- | --- | --- | --- | --- | --- | --- |
| Left<br>mPFC | Right<br>mPFC | 3.00 | 1790.10 | 1.58 | 1.18 | Within default mode<br>network | Homotopic interhemispheric |
| Right<br>mPFC | Left<br>mPFC | 3.00 | 1790.10 | 1.58 | 1.18 | Within default mode<br>network | Homotopic interhemispheric |
| Left<br>DI | Left<br>GI | 2.96 | 1046.70 | 1.30 | 0.31 | No | Intrahemispheric left |
| Left<br>GI | Left<br>DI | 2.89 | 1046.70 | 1.30 | 0.31 | No | Intrahemispheric left |
| Left<br>LPtA | Left<br>S1Tr | 3.00 | 629.20 | 1.28 | 0.97 | Sensorimotor network to<br>another network | Intrahemispheric left |
| Left<br>S1Tr | Left<br>LPtA | 3.00 | 629.20 | 1.28 | 0.97 | Sensorimotor network to<br>another network | Intrahemispheric left |
| Left<br>V2L | Left<br>V1B | 3.00 | 1474.60 | 1.28 | 1.43 | No | Intrahemispheric left |
| Left<br>M1 | Left<br>M2 | 2.96 | 3787.60 | 1.28 | 1.27 | Within sensorimotor<br>network | Intrahemispheric left |
| Left<br>M2 | Left<br>M1 | 3.76 | 3787.60 | 1.28 | 1.27 | Within sensorimotor<br>network | Intrahemispheric left |
| Right<br>LPtA | Right<br>S1Tr | 3.00 | 533.70 | 1.27 | 0.98 | Sensorimotor network to<br>another network | Intrahemispheric right |
| Right<br>S1Tr | Right<br>LPtA | 3.00 | 533.70 | 1.27 | 0.98 | Sensorimotor network to<br>another network | Intrahemispheric right |
| Left<br>GI | Left<br>S2 | 3.72 | 1075.40 | 1.25 | 1.64 | Sensorimotor network to<br>another network | Intrahemispheric left |
| Left<br>S2 | Left<br>GI | 3.62 | 1075.40 | 1.25 | 1.64 | Sensorimotor network to<br>another network | Intrahemispheric left |
| Right<br>M1 | Right<br>M2 | 2.96 | 2958.90 | 1.22 | 1.27 | Within sensorimotor<br>network | Intrahemispheric right |
| Right<br>M2 | Right<br>M1 | 3.76 | 2958.90 | 1.22 | 1.27 | Within sensorimotor<br>network | Intrahemispheric right |
| Right<br>GI | Right<br>S2 | 3.72 | 1261.10 | 1.22 | 1.63 | Sensorimotor network to<br>another network | Intrahemispheric right |
| Right<br>S2 | Right<br>GI | 3.62 | 1261.10 | 1.22 | 1.63 | Sensorimotor network to<br>another network | Intrahemispheric right |
| Right<br>DI | Right<br>GI | 2.96 | 1200.00 | 1.21 | 0.31 | No | Intrahemispheric right |
| Right<br>GI | Right<br>DI | 2.89 | 1200.00 | 1.21 | 0.31 | No | Intrahemispheric right |
| Left<br>M1 | Left<br>S1FL | 2.88 | 1189.60 | 1.20 | 1.90 | Within sensorimotor<br>network | Intrahemispheric left |
| Left<br>S1FL | Left<br>M1 | 2.86 | 1189.60 | 1.20 | 1.90 | Within sensorimotor<br>network | Intrahemispheric left |
| Right<br>M1 | Right<br>S1FL | 2.88 | 1077.70 | 1.20 | 1.91 | Within sensorimotor<br>network | Intrahemispheric right |
| Right<br>S1FL | Right<br>M1 | 2.86 | 1077.70 | 1.20 | 1.91 | Within sensorimotor<br>network | Intrahemispheric right |
| Left<br>AuD | Left AuI | 3.00 | 1098.90 | 1.18 | 0.78 | Within default mode<br>network | Intrahemispheric left |
| Right<br>Cg1 | Left<br>mPFC | 3.00 | 260.00 | 1.18 | 2.87 | Within default mode<br>network | Heterotopic<br>interhemispheric |
| Left<br>DI | Left AID | 3.00 | 851.00 | 1.16 | 3.05 | No | Intrahemispheric left |

Seed and target regions were determined from the NeuroVIISAS tracer database. AID: agranular insular cortex dorsal part; AuI: primary auditory cortex; AuD: secondary auditory cortex dorsal area; Cg1: cingulate cortex area 1; DI: dysgranular insular cortex; GI: granular insular cortex; LPtA: lateral parietal association cortex; M1: primary motor cortex; M2: secondary motor cortex; mPFC: medial prefrontal cortex; S1FL: primary somatosensory cortex forelimb region; S1Tr: primary somatosensory cortex trunk region; S2: secondary somatosensory cortex; V1B: primary visual cortex binocular area; V2L: secondary visual cortex lateral area.

Table 3 shows the connections that we identified as belonging to the 25% weakest structural and functional connections. The average length of the weakest structural and functional connections was 10.84 ± 2.09 mm. The identified connections included 30% heterotopic interhemispheric and 70% intrahemispheric (left: 44%, right: 26 %) connections, and were mainly between frontal cortices, parahippocampal areas and the retrosplenial cortex. Sixty-one percent of the connections connected the default mode network with another functional network.

**Table 3:**
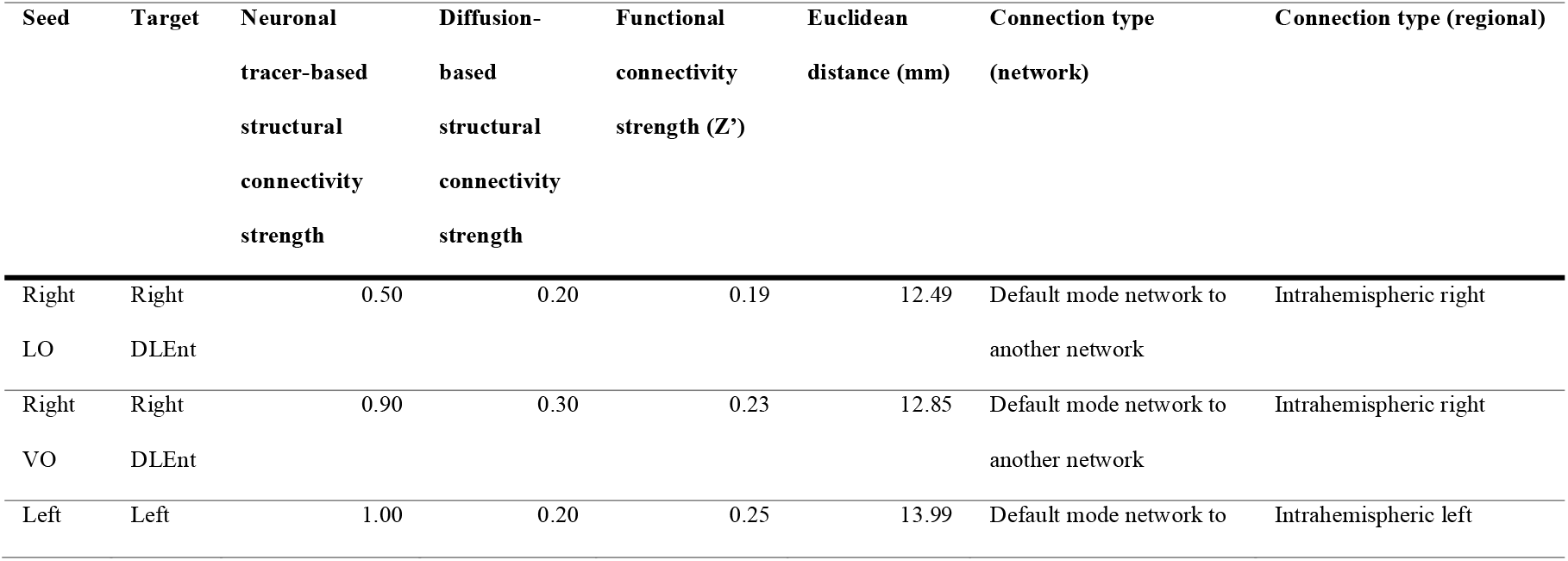
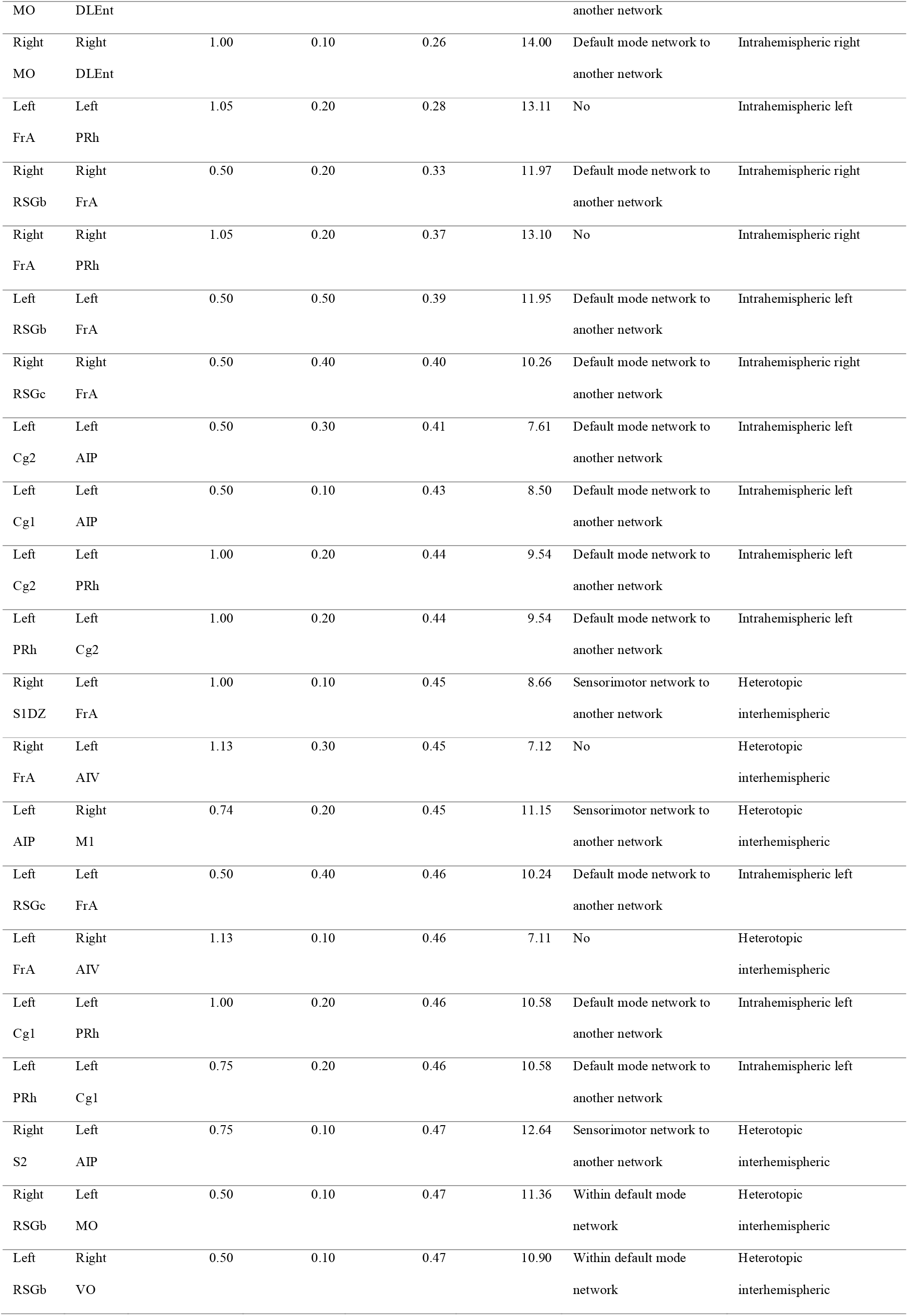
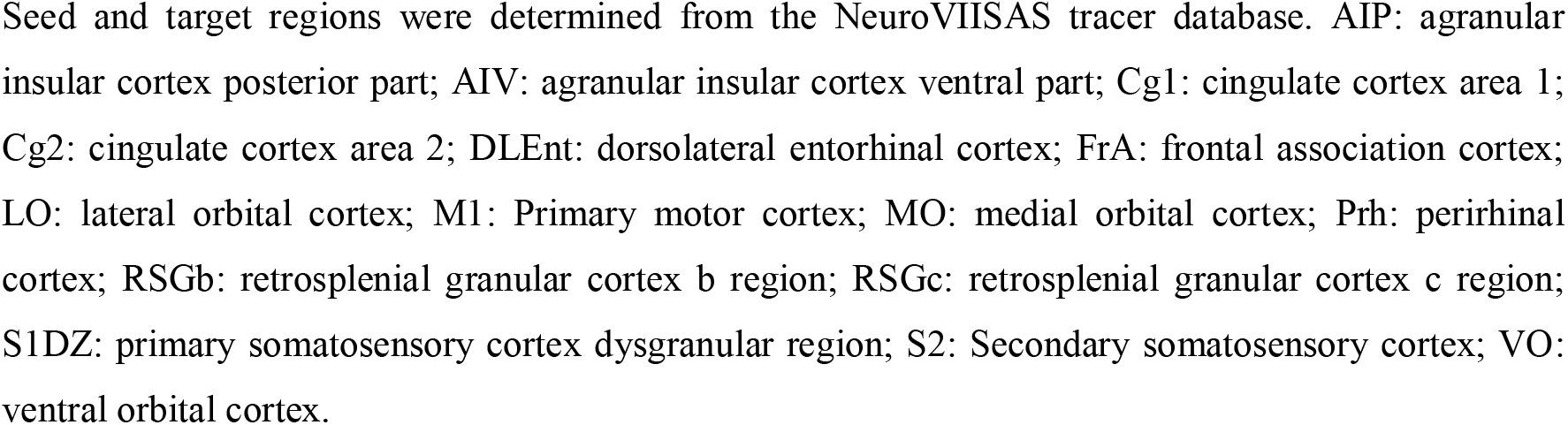
Characteristics of cortical connections in the rat brain with weak meso- and macroscale structural connectivity and weak functional connectivity.

Table 4 shows connections with strong structural but weak functional connectivity. All these connections were intrahemispheric (50% right; 50% left), of which the average Euclidean distance between regions was 3.23 ± 1.64 mm. Many of these connections were between parahippocampal areas and the insular cortex or auditory cortex, and within the insular cortex.

**Table 4:**
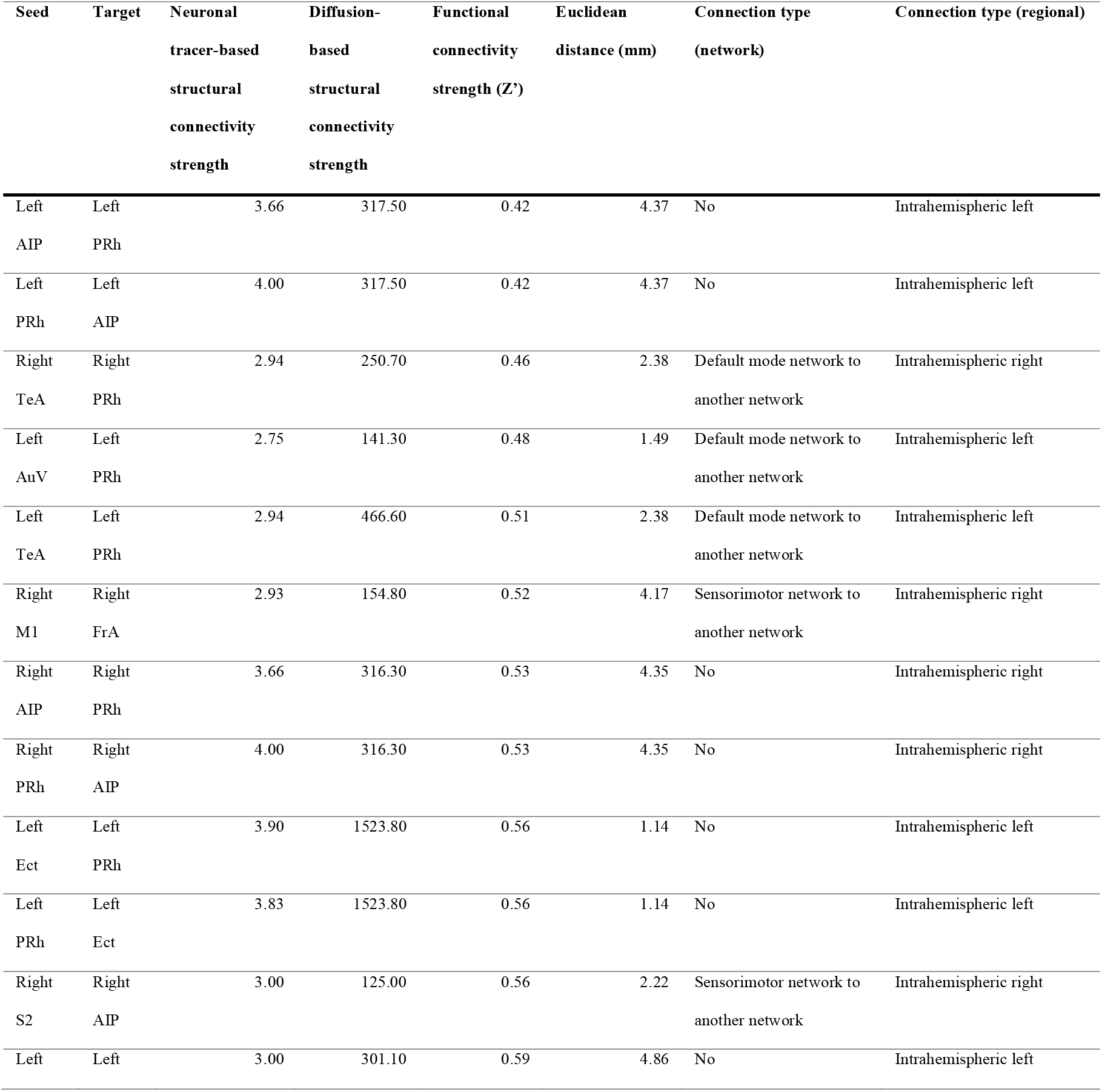
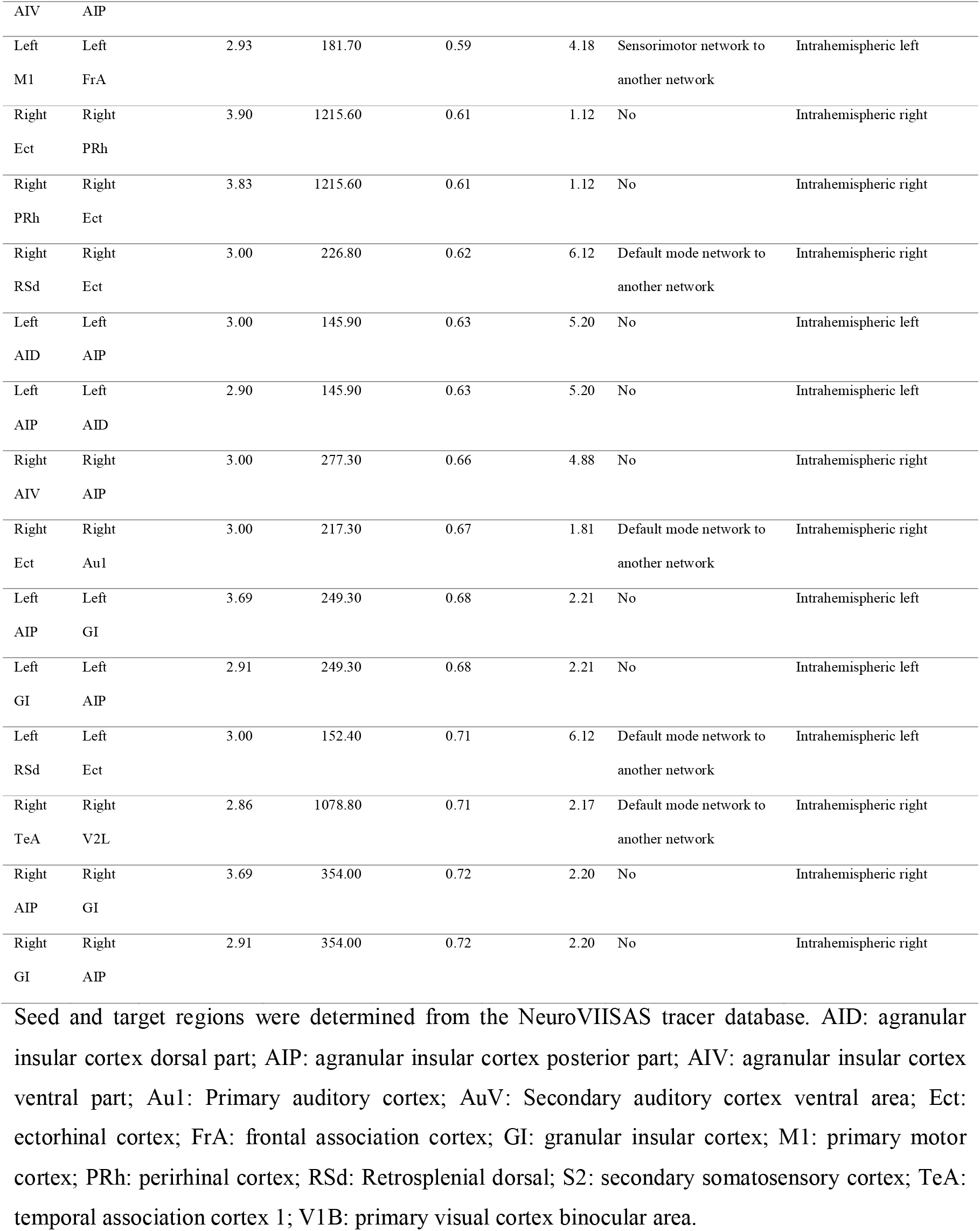
Characteristics of cortical connections in the rat brain with strong meso- and macroscale structural connectivity and weak functional connectivity.

The connections belonging to 25% strongest functional but 25% weakest structural connections are shown in Table 5. The average Euclidean length of the connections was 8.67 ± 1.78 mm and 35% were heterotopic interhemispheric, all of which were part of the sensorimotor network. Fifty-two percent of the connections resided within the sensorimotor network or between the sensorimotor network and another network, of which 42% connected the sensorimotor with the default mode network, and 43% of the connections was between the default mode network and another functional network.

**Table 5:**
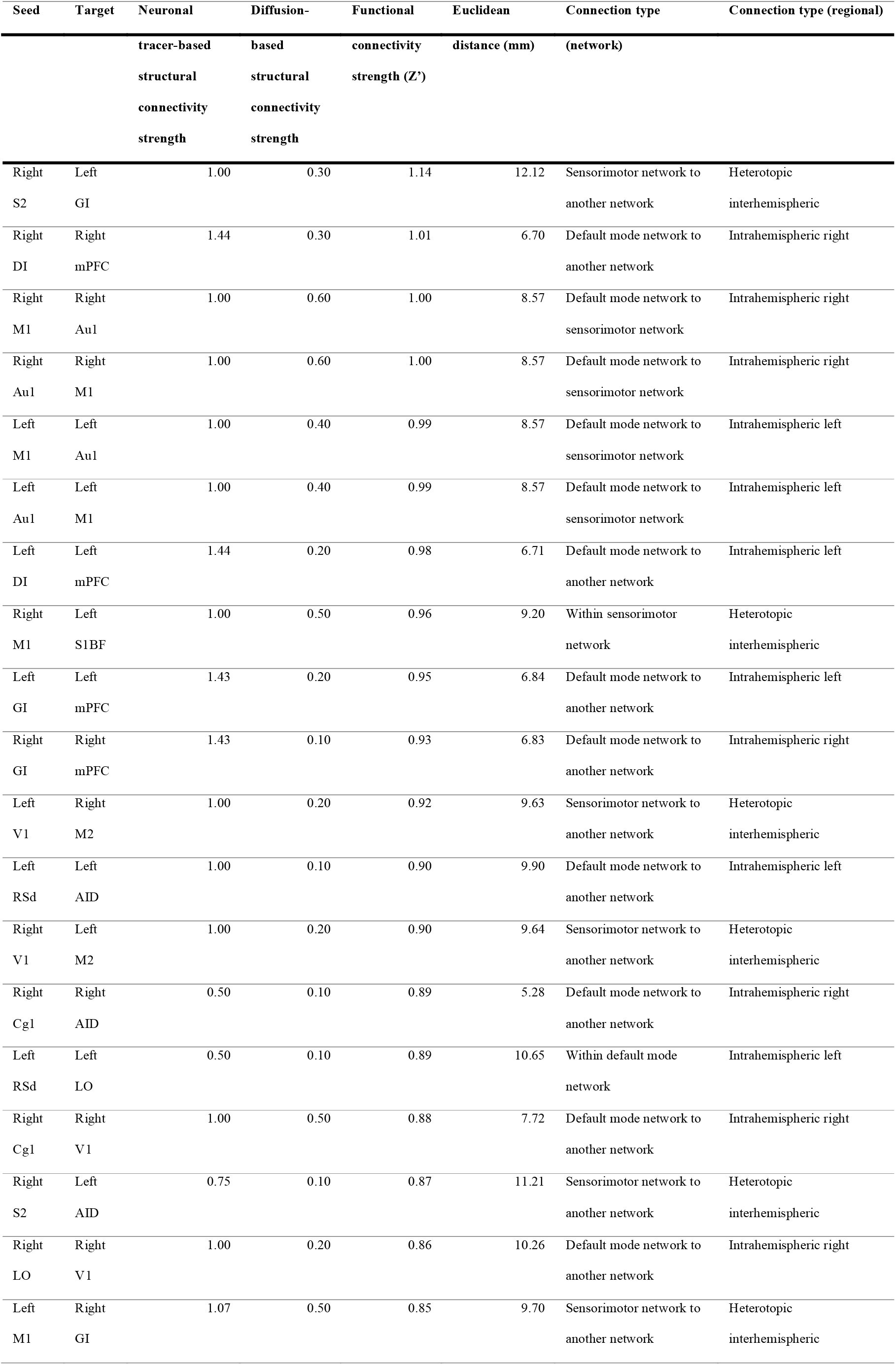
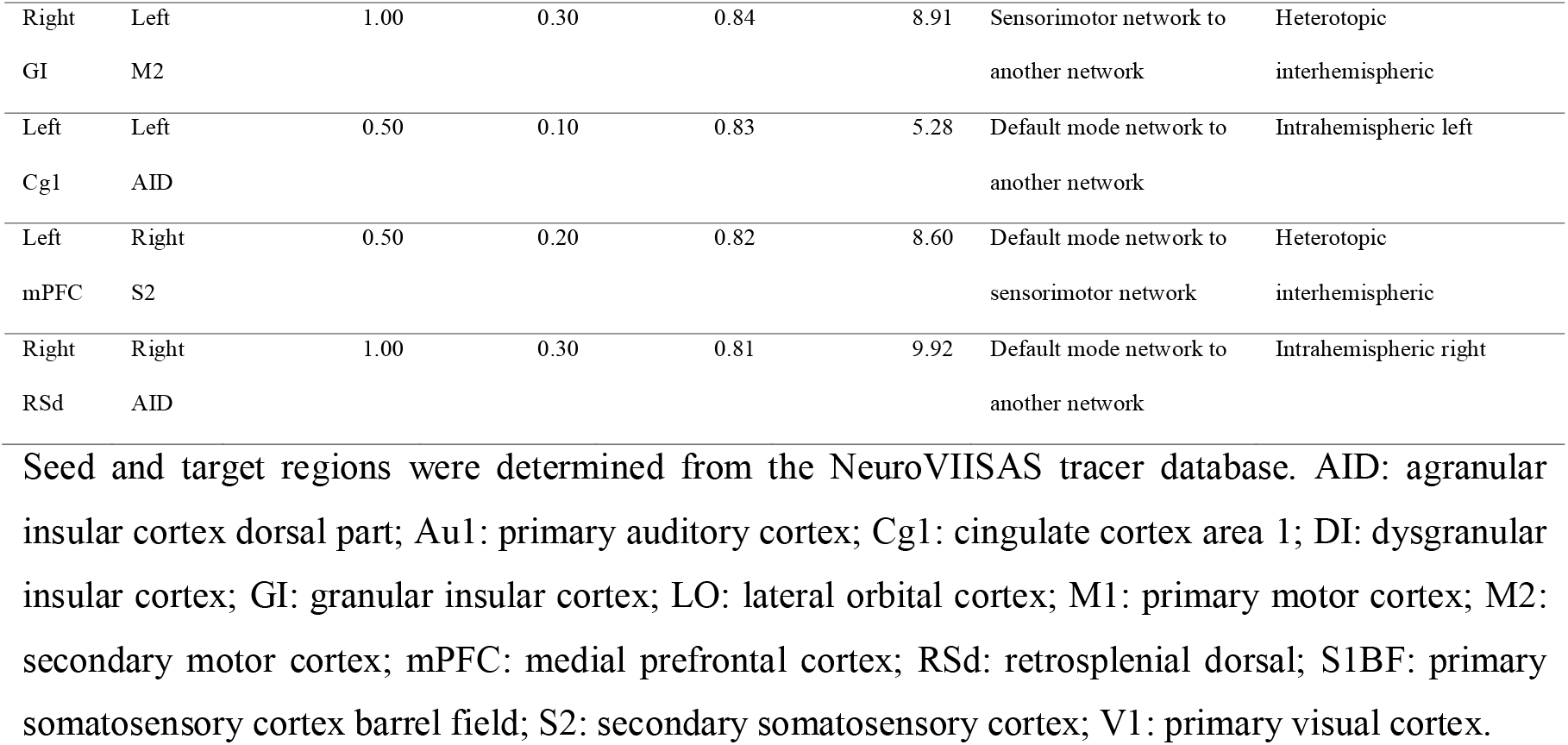
Characteristics of cortical connections in the rat brain with weak meso- and macroscale structural connectivity and strong functional connectivity.

### Global correlation between structural and functional connectivity depends on method and scale

Functional connectivity strength was positively correlated with diffusion-based structural connectivity strength in cortical connections (ρ=0.41; p<0.0001; Figure 4a). For the same cortical connections, functional connectivity strength did not correlate with neuronal tracer-based structural connectivity strength (ρ=0.04, p=0.14; Figure 4b).

**Figure 4:**
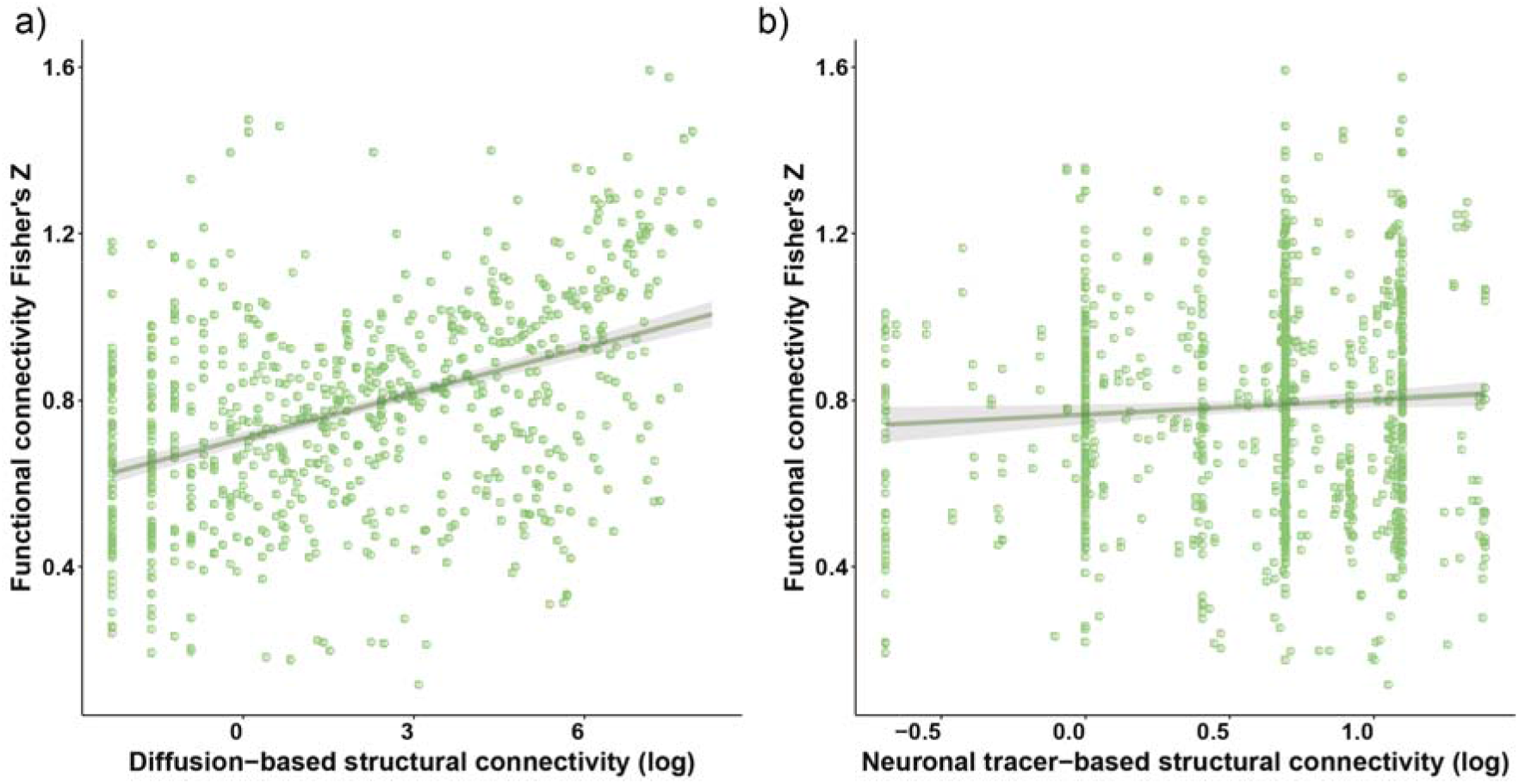
Whole-brain structure-function relationships at the structural macro-scale (diffusion-based structural connectivity) and meso-scale (neuronal tracer-based structural connectivity). Functional connectivity strength is plotted as the Fisher’s Z-transformed correlation coefficient versus the log-transformed diffusion-based (a) or neuronal tracer-based structural connectivity strength (b). Individual connections are plotted as green dots. The structure-function relationship is shown as a linear fit, with shading representing the 95% confidence intervals of the fit.

## Discussion

Our study on the rat brain shows that cortical brain networks are characterized by functional connectivity strengths, as measured with resting-state fMRI, that partly associate with macro-scale diffusion-based structural connectivity strength but not associate with meso-scale neuronal tracer-based structural connectivity strength. At a more local level we found that strong functional connectivity in the sensorimotor and default mode network matched with strong structural connectivity of intrahemispheric connections but was accompanied by weak structural connectivity of interhemispheric connections.

### Distinct global structure-function relationships across different hierarchical levels of structural connectivity

The partial positive correspondence between functional connectivity and diffusion-based structural connectivity strength in the rat brain is in line with structure-function relationships found in humans (Straathof et al. 2019). However, we did not find a correlation between functional connectivity and meso-scale neuronal tracer-based structural connectivity strength. One previous study investigated this relationship at the meso-scale in rats and reported a positive structure-function correlation (r=0.48) (Díaz-Parra et al. 2017). However, this study did not include essential interhemispheric connections. Interhemispheric connections are known to have lower structurefunction relationships (Shen et al. 2012), which may be explained by long inter-regional distances, sparser interhemispheric connectivity or involvement of polysynaptic or indirect connections (O’Reilly et al. 2013). Distinct structure-function relationships at the structural macro- and meso-scale have already been demonstrated in a study combining datasets in humans (functional and diffusion-based structural connectivity) and macaques (neuronal tracer-based structural connectivity) (Reid et al. 2016). However, the authors could not disentangle whether these distinct relationships were due to species differences or due to different measures of structural connectivity. Since we compared all three measures in the same species, (dis)agreement between structural and functional connectivity most likely reflects topological differences in the structure-function relationship across different hierarchical levels.

Beside being measurements at different hierarchical levels, another important difference between macro-scale diffusion-based and meso-scale neuronal tracer-based structural connectivity is the directionality information available in the data. Whereas diffusion-based structural connectivity does not provide directionality information, meaning that all connections are considered to be fully reciprocal, neuronal tracer-based structural connectivity does provide this directionality information. Since resting-state functional connectivity is also directionless, the correlation of functional connectivity with diffusion-based structural connectivity may be higher than with neuronal tracer-based structural connectivity. In addition, the correlation between functional connectivity and diffusion-based structural connectivity may also be explained by the fact that both connectivity measures are determined with the same measurement tool, i.e. MRI.

### Strong functional connectivity in robust resting-state networks is supported by strong short-range intrahemispheric connections

The sensorimotor and default mode network are robustly established resting-state networks in the rodent brain (Pawela et al. 2008; Sierakowiak et al. 2015), which was corroborated by our finding of strong functional connectivity in or between these networks. We also observed strong short-range intrahemispheric structural connections at meso- and macro-scale in these networks. Strong reciprocal structural connections have previously been shown between ipsilateral sensorimotor cortices, measured with neuronal tracers (Miyashita et al. 1994; Hoffer et al. 2003; Rocco-Donovan et al. 2011), and between default mode network regions, measured with diffusion MRI (Greicius et al. 2009; Horn et al. 2014). In comparison, in the current study we found that heterotopic interhemispheric structural connections in the sensorimotor network and long-range intrahemispheric structural connections between the default mode network and other functional networks were weak at both the macro- and the meso-scale. Since both connection types were between areas located far apart from each other, this observation may reflect the difficulties of diffusion-based tractography to reconstruct long-distance connections (Reveley et al. 2015), and the distance-dependence of neuronal tracer-based structural connectivity strength (Ercsey-Ravasz et al. 2013).Our data point out that the distance-dependence of structural connectivity strength, as determined from diffusion MRI or neuronal tracing, influences measurements of structure-function relationships. This should be taken into account in studies on the relation between structural and functional connectivity. However, weak heterotopic interhemispheric connectivity may also reflect the smaller role these connections play in functional brain organization as compared to homotopic interhemispheric connections (Deco et al. 2014; Messé et al. 2014). Interestingly, strong functional connectivity in homotopic interhemispheric connections within the sensorimotor network was not accompanied by strong structural connectivity, despite the presence of a large bundle of neuronal fibers, i.e. the corpus callosum, connecting the two hemispheres. This may be a result of our approach of only including connections that exhibit macro- and meso-scale structural connectivity. Homotopic interhemispheric connections in the sensorimotor network were included in the 25% strongest meso-scale neuronal tracer-based structural network, but not in the 25% strongest macro-scale diffusion-based structural network. Therefore, we limit our conclusions to connections with matching macro- and meso-scale structural connectivity, while other structure-function relationships may exist in connections where macro- and meso-scale structural connectivity do not match.

### Implications of different structure-function relationships across the brain in health and disease

We have shown that different structure-function relationships exist in different cortical connections of the rat brain, in line with a previous study reporting that 25% of valid structural connections are very weak functional connections (Lee and Xue 2018). Different structure-function relationships can have implications for brain functioning and behavior. Healthy brain functioning relies on a balance between segregation and integration of neuronal communications (Tononi et al. 1994; Fox and Friston 2012). Structure-function relationships have been shown to be stronger when functional networks are in an integrated state, compared to a segregated state (Fukushima et al. 2018). In another study, white matter integrity was associated with BOLD signal complexity in local connections (structure-function agreement) but not in distributed connections (structure-function disagreement) (Mcdonough and Siegel 2018). This suggests that information integration relies on a strong structure-function relationship, whereas weak structure-function relationships are implied in segregation.

Next to the implication of structure-function relationships on healthy brain functioning, structure-function relationships may (partly) determine the functional effects of structural damage to the brain. Intuitively, it may be deduced that structural damage to connections with strong structurefunction relationships will have severer functional consequences than structural damage to connections with weak structure-function relationships. Novel algorithms may enable us to predict the functional effects of specific structural damage (Meier et al. 2016). Alterations and preservations of structural and functional connectivity in human patients, and in animal models of neurological and psychiatric diseases, can provide insights into the impact of structure-function couplings on outcome. For example, after stroke significant changes in structural and functional connectivity have been measured in the remaining intact sensorimotor network in rodents (van Meer et al. 2010, 2012; Schmitt et al. 2017) and humans (Carter et al. 2010; Radlinska et al. 2012; Grefkes and Fink 2014). Chronically after experimental stroke in rats, structural and functional connectivity changes were related intrahemispherically –on the side of the stroke lesion–while this was not evident for interhemispheric connections (van Meer et al. 2010). This may be explained by a stronger structurefunction agreement in intrahemispheric sensorimotor connections as compared to interhemispheric sensorimotor connections, as we found in the current study.

A strength of the current study is the inclusion of three different measures of connectivity within a single species. Comparing functional connectivity against macro-scale diffusion-based as well as meso-scale neuronal tracer-based structural connectivity in rats enabled the investigation of structure-function relationships across hierarchical levels. In addition, by including both diffusion- and neuronal tracer-based structural connectivity measures, we could avoid inclusion of false positives that are often present in diffusion-based structural networks (Calabrese et al. 2015; Chen et al. 2015; Maier-Hein et al. 2017; Sinke et al. 2018). A reliable structural network of the rat brain was created by only selecting connections present in both diffusion- and neuronal tracer-based structural networks. The relationship between diffusion-based structural connectivity and resting-state functional connectivity may have been higher when both measures would have been acquired in the same rat. However, neuronal tracer-based structural connectivity was acquired from many different groups of rats. Therefore, we also measured diffusion-based structural connectivity in a separate group of rats, to prevent inappropriate comparison with potentially higher within-subject correlations. A limitation could be the restriction of our assessments to monosynaptic connections. In addition, resting-state functional connectivity was determined under anesthesia, which influences functional connectivity measures (Paasonen et al. 2018) and possibly affects the structure-function relationship.

In conclusion, we demonstrated a correlation between functional connectivity and diffusion-based structural connectivity, but no correlation between functional connectivity and neuronal tracer-based structural connectivity in the rat cortex. These distinct structure-function relationships may be due to different hierarchical levels of measurement or directionality information available in the data. In addition, the structure-function relationship varies across cortical regions in the rat brain. Characteristics of the used techniques, such as distance-dependency, affect where structural and functional networks (dis)agree. Conclusions about connectivity based on a single technique may therefore be biased. This shows the importance of combining different complementary measures of connectivity at distinct hierarchical levels to accurately determine connectivity across networks in the healthy and diseased brain.

## Acknowledgement

This is a pre-print of an article published in Scientific Reports. The final authenticated version is available online at: https://doi.org/10.1038/s41598-019-56834-9.

